# Beyond Uplands: Integrating Riparian Areas into Sagebrush Conservation Design for a More Resilient West

**DOI:** 10.64898/2025.12.29.696625

**Authors:** Kristopher R. Mueller, Scott L. Morford, Jeremy D. Maestas, John S. Kimball, Andrew C. Olsen, David J.A. Wood, David E. Naugle

**Affiliations:** Numerical Terradynamic Simulation Group, University of Montana, Missoula, MT 59812, USA; United States Department of Agriculture, Natural Resources Conservation Service, Bend, OR, 97702, USA; Intermountain West Joint Venture, Missoula, MT, 59801, USA; United States Department of the Interior, Bureau of Land Management, Billings, MT, 59101, USA; W. A. Franke College of Forestry and Conservation, University of Montana, Missoula, MT 59812, USA

## Abstract

Riparian and mesic habitats are essential biological hotspots in the American West, supporting 80% of wildlife, yet they are often assessed in isolation from surrounding uplands. To address this, we developed a framework that complements the Sagebrush Conservation Design to incorporate these habitats into broader landscape conservation planning. By analyzing 40 years of riparian vegetation data alongside sagebrush ecosystems, we classified watersheds into three actionable management tiers based on water availability and vegetation health. This prioritization is operationalized through the Mesic Analysis Platform, an interactive tool that identifies stream reaches with floodplain disconnection as key targets for restoration. Notably, the greatest restoration potential exists on private lands, where 63–65% of identified mesic habitats are found. This framework provides a practical strategy to integrate social feasibility with ecological priority, fostering the multi-jurisdictional partnerships vital to connecting water resources with the broader sagebrush ecosystem.

## Introduction

Riparian ecosystems, including river corridors, valley floodplains, and wetlands–termed mesic habitats–are among Earth’s most biologically diverse yet imperiled landscapes (Dudgeon et al., 2006). Despite occupying only a small fraction of the land, these areas deliver a disproportionately large suite of ecosystem services. They support high biodiversity, regulate water quality, and provide flood mitigation (Riis et al., 2020). Globally, these systems have declined, with estimates suggesting that up to 90% of floodplains in Europe and North America are now functionally lost (Schneider et al., 2017). This widespread degradation is largely driven by multiple, compounding threats, including hydrologic alteration from dam construction and water diversions, which disrupt natural flow regimes and diminish ecologically important flood pulses (Nilsson & Berggren, 2000; Schneider et al., 2017). This is exacerbated by land-use change, which fragments habitat through cultivation and urbanization (Patten, 1998), and by climate change, which heightens vulnerability, particularly in arid regions, by increasing water stress and altering disturbance regimes (Theobald et al., 2010). The pervasive loss and degradation of these water-dependent ecosystems demand strategic, large-scale conservation frameworks to conserve and restore their remaining functions.

### The Challenge of Mesic Habitat Conservation

In the arid and semi-arid sagebrush biome of the western U.S., the importance of water-dependent ecosystems is amplified by water scarcity, making them highly vulnerable to degradation. These moisture-dependent areas sustain green vegetation during dry periods and provide essential resources that enable wildlife to persist in adjacent uplands (Fig. 1). However, mesic ecosystems have experienced significant changes. These alterations have mostly damaged hydrologic functions and plant structures, leading to widespread ecosystem decline. In the past, this degradation was a result of widespread beaver removal from the landscape, overgrazing by cattle, water diversions, and land conversion, but has since shifted to also include the expansion of invasive species and climate change, enhancing degradation (Poff et al., 2011). These complex, large-scale challenges make it difficult for practitioners to prioritize limited restoration resources effectively. Successful management and restoration of mesic habitats require clear prioritization frameworks and accessible tools. Both scientists and managers identify the need for accessible tools and the prioritization of ecosystem services at multiple spatial scales as a top challenge for riparian conservation globally (Rodríguez-González et al., 2022). Existing restoration frameworks often fail to provide the landscape-scale context, particularly the critical interdependence between mesic habitats and adjacent upland habitat quality required for strategic conservation decisions across western U.S. rangelands. This gap in landscape-scale planning is a significant obstacle to conservation efforts.

**Figure 1.**
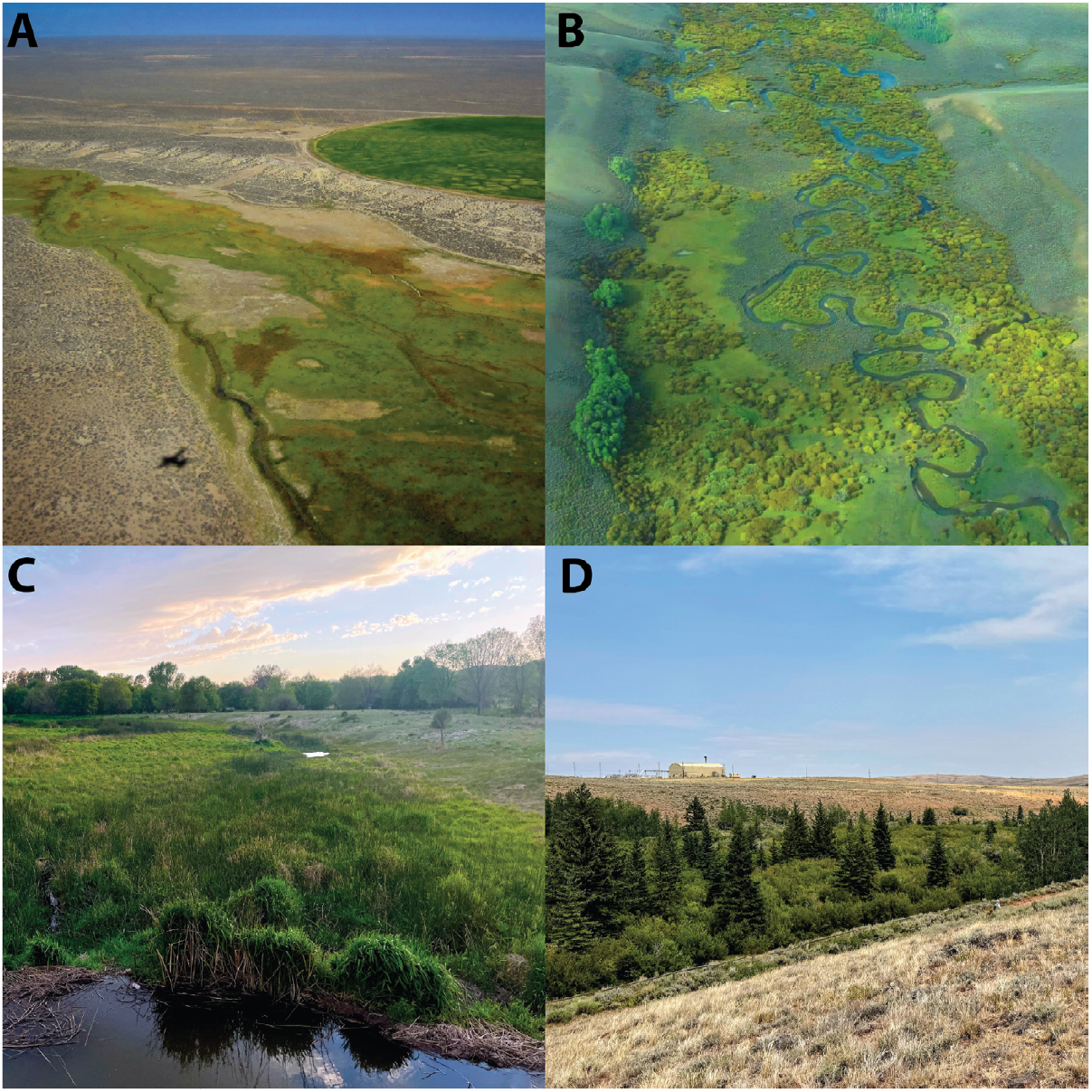
Examples of mesic habitats in valley bottoms of the western United States. (A) A mesic floodplain adjacent to an upland area with center-pivot irrigation. (B) A meandering stream supporting a well-developed riparian corridor. (C) A beaver dam impounds water and increases productivity within a stream reach. (D) A riparian valley bottom contrasts with the surrounding arid uplands.

### Mesic Area Prioritization as a Compliment to the Sagebrush Conservation Design

The Sagebrush Conservation Design (SCD) is one of the most comprehensive tools for conservation planning in the sagebrush biome, providing a strategic approach to management across 175 million acres of western U.S. sagebrush rangelands (Doherty et al., 2022). This spatially targeted, proactive, and collaborative framework is an effective tool for reversing sagebrush decline throughout the western U.S (Mozelewski et al., 2024; sagebrushconservation.org). It uses a ‘Defend and Grow the Core’ strategy, focusing on protecting and growing intact Core Sagebrush Areas (CSAs) and restoring Growth Opportunity Areas (GOAs). While effective for conserving extensive upland sagebrush habitats, the SCD does not include mesic habitats in its scope, creating a significant gap in conservation planning. Mesic habitats comprise only 2% of arid western landscapes but support up to 80% of land-based wildlife, providing essential cover and food resources during dry conditions (Maestas et al., 2023; Naiman et al., 1993; Thomas et al., 1979). These habitats maintain vegetative productivity through steady access to shallow groundwater, sustaining wildlife populations that require both mesic and upland resources (Kolarik et al., 2023; Manning et al., 2020; Noy-Meir, 1973; Shrestha et al., 2024). Accounting for these mesic habitats would further enhance the SCD’s ecosystem conservation approach, ensuring protection of the critical landscapes essential for wildlife.

In this paper, we argue that prioritizing mesic areas is essential for maintaining functional sagebrush ecosystems and their wildlife populations. We present a complementary framework and tool that identifies mesic conservation opportunities across the entire biome, then highlights those near intact sagebrush uplands where protection and restoration will yield the greatest benefits for ecosystem function and wildlife persistence. Our approach further provides practitioners with actionable guidance for integrating mesic conservation into existing sagebrush management strategies.

### Leveraging Remote Sensing for Targeted Conservation

Advances in remote sensing now enable landscape-scale assessment of mesic conservation and restoration opportunities. While existing tools map vegetation cover (Allred et al., 2021) and late-season mesic vegetation extent (Donnelly et al., 2016) across western rangelands, these data are often too coarse for precise conservation targeting. Our approach uses these established datasets and builds upon them in a new interactive planning tool called the Mesic Analysis Platform (MAP). Although managers may depend on additional information to identify priority conservation opportunities within the sagebrush biome, MAP offers an inventory of mesic habitats that can identify individual stream reaches at a scale suitable for guiding management actions. This enables practitioners to pinpoint areas where field assessments will be most beneficial and where management efforts may have the highest potential for success. By analyzing productivity patterns along stream networks, we developed a framework to identify mesic habitats with potentially degraded floodplains and inconsistent water availability. When these areas are adjacent to intact, water-rich habitats, they signal a hydrologic disconnection that practitioners can then verify and address. This method converts landscape data into practical, reach-scale targets, streamlining the process of identifying site-specific interventions.

### Restoration Approaches and Context

Once mesic habitat priorities are identified, restoration methods must align with the scale and context of each system. A majority of mesic resources, or vegetation in mesic habitats, occur within large-scale primarily irrigated floodplains, where agricultural practices on privately owned lands sustain much of the available mesic habitats (Donnelly et al., 2016, 2024). While these working landscapes are critical for biodiversity and habitats, direct restoration of larger river floodplains is often cost and scale-prohibitive, placing them outside the scope of this paper. In contrast, smaller order streams and degraded headwaters represent systems where low-tech restoration practices are highly effective. In these settings, Low-Tech Process-Based Restoration (LTPBR) has become a particularly cost-effective approach for water-limited landscapes (Wheaton et al., 2019). LTPBR deploys simple structures to restore natural stream functions, reconnect floodplains, and raise water tables, which helps promote mesic vegetation by increasing soil moisture in smaller stream reaches (Pollock et al., 2014; Shrestha et al., 2024; Zeedyk & Clothier, 2014). These techniques, alongside grazing management practices, have demonstrated durability and effectiveness. For example, mesic areas with Zeedyk structures (hand-built rock and soil structures in stream) showed higher vegetation productivity and resilience to drought 20 years following restoration, as observed through satellite imagery (Silverman et al., 2019).

To enhance and add more nuance to the current SCD framework, we developed a complementary framework and tool specifically for integrating mesic habitats. Our framework identifies intact, water-rich areas for protection and potentially water-limited sites with restoration potential across the sagebrush biome, prioritizing areas near intact sagebrush uplands where conservation actions can enhance functional mesic habitats and maximize broader ecosystem interconnectedness. By incorporating mesic habitats into conservation planning, we reveal additional targeting opportunities to create a more connected and resilient sagebrush biome. Specifically, we:

1. Analyzed the spatial relationships between persistent mesic vegetation and intact sagebrush uplands across the western U.S. using 40 years of vegetation productivity data.
2. Developed a tiered mesic conservation actions framework and online tool to identify watersheds with the highest mesic conservation and restoration potential in the sagebrush biome.

## A Critical Disconnect: Mesic Hotspots and Sagebrush Cores Rarely Overlap

The effectiveness of ecosystem conservation depends on understanding how mesic habitats spatially relate to intact sagebrush uplands prioritized by the SCD. To evaluate whether current conservation adequately addresses ecosystem-wide habitat needs, we analyzed 40 years of vegetation data to map productive mesic habitats in relation to intact sagebrush uplands across the biome.

### Methods for Spatial Analysis

We defined mesic habitats within riparian floodplains using valley bottom extents from the Valley Bottom Extraction Tool (VBET), which identifies floodplain valleys’ potential for water inundation under normal hydrologic conditions based on 10-meter elevation models and National Hydrography Dataset (NHD) delineated stream channels (Gilbert et al., 2016). Valley bottom extents are divided into smaller sections, described here as “reaches”, or Discrete Geographic Objects (DGOs) in Gilbert et al. (2016). VBET data were collected through the Riverscapes Data Exchange (https://data.riverscapes.net/pt/vbet), and the extents with segmented reaches were imported into Google Earth Engine for analysis (Gorelick et al., 2017). Our analysis focused on the sagebrush biome to align with the SCD spatial extent. Additionally, the region’s dry summer conditions create a strong spectral contrast between mesic and upland vegetation, aiding classification through optical-infrared remote sensing (Weier & Herring, 2000). Satellite imagery from the Landsat 5, 7, 8, and 9 series was used to identify productive mesic vegetation across the biome at a 30-meter spatial resolution. We used the Landsat-derived Normalized Difference Vegetation Index (NDVI) with a mean threshold of ≥ 0.3 during the late growing season (July 15 to September 30) to identify productive mesic vegetation during the most water-limited time of the year (Donnelly et al., 2016, 2018; Lundblad et al., 2022).

We then calculated productive vegetation persistence for each pixel from 1984 to 2024 (excluding 2012 due to limited Landsat imagery coverage), as a proxy for water availability in dryland systems (Shrestha et al., 2024). Pixels were classified into three persistence categories based on the number of years with identified productive vegetation over the time series: high persistence (productive ≥ 75% of years), moderate persistence (productive ≥ 25% and < 75% of years), low persistence (productive 1 - 24% of years), and no persistence (no identified late growing productive vegetation). Classified pixels were then filtered to include only pixels within valley bottom extents, excluding cultivated and urban pixels. Intact sagebrush uplands were defined as either CSAs or GOAs using the 2020 SCD model with a five-kilometer geographic buffer, based on proximal use of mesic habitats by wildlife (Donnelly et al., 2016). The total pixel area of each persistence class and nearby intact sagebrush uplands was then calculated and summarized within valley bottom extents. The most recent Protected Area database (USGS, 2024; https://www.usgs.gov/programs/gap-analysis-project/science/pad-us-data-overview) was used to delineate public versus private lands.

### Spatial Mismatch Between Floodplain Mesic Habitats and Intact Sagebrush Uplands

Our analysis reveals a stark spatial disconnect: the most persistent mesic habitats in riparian floodplains rarely overlap with intact sagebrush uplands across the biome. High-persistence mesic habitats concentrate in forested ecosystems and montane foothills, extending into rangelands only where water resources are likely sufficient (Fig. 2A). In contrast, intact sagebrush uplands occur in open rangelands with scarce water and mesic habitats (Fig. 2B). However, these upland areas may contain localized, drought-resistant vegetation that provides critical forage throughout the year. This spatial separation is particularly evident in eastern Oregon and central Wyoming. Critically, less than 10% of identified high-persistence mesic habitats are within 5 kilometers of intact sagebrush uplands (Table 1), severely limiting the availability of complementary habitats essential for wildlife persistence during dry periods.

**Table 1.**
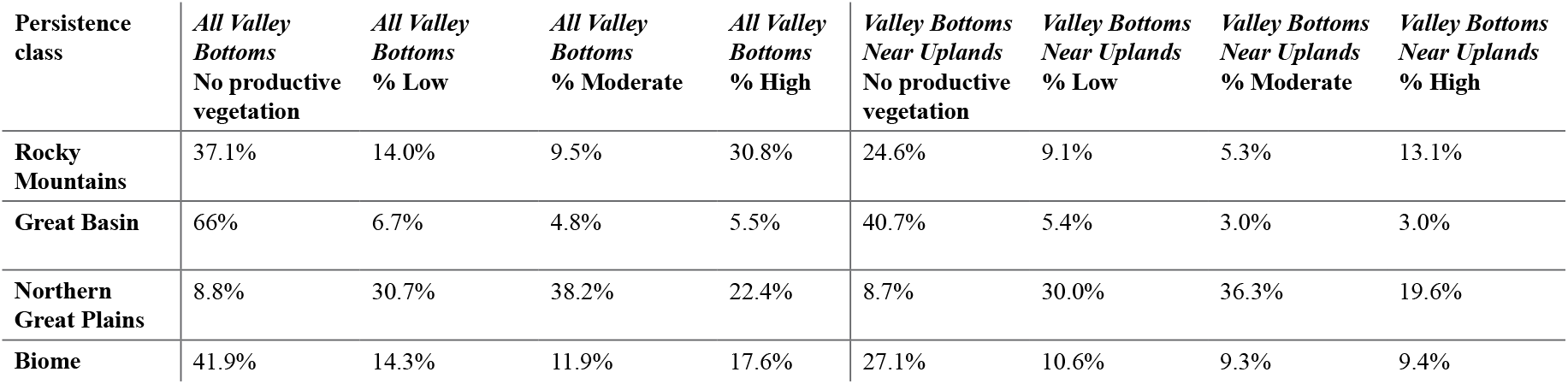
Proportional area of mesic habitat persistence classes within valley bottoms across the sagebrush biome. Values represent the percentage of the total valley bottom area, comparing all valley bottoms versus those within 5 km of intact uplands.

**Figure 2.**
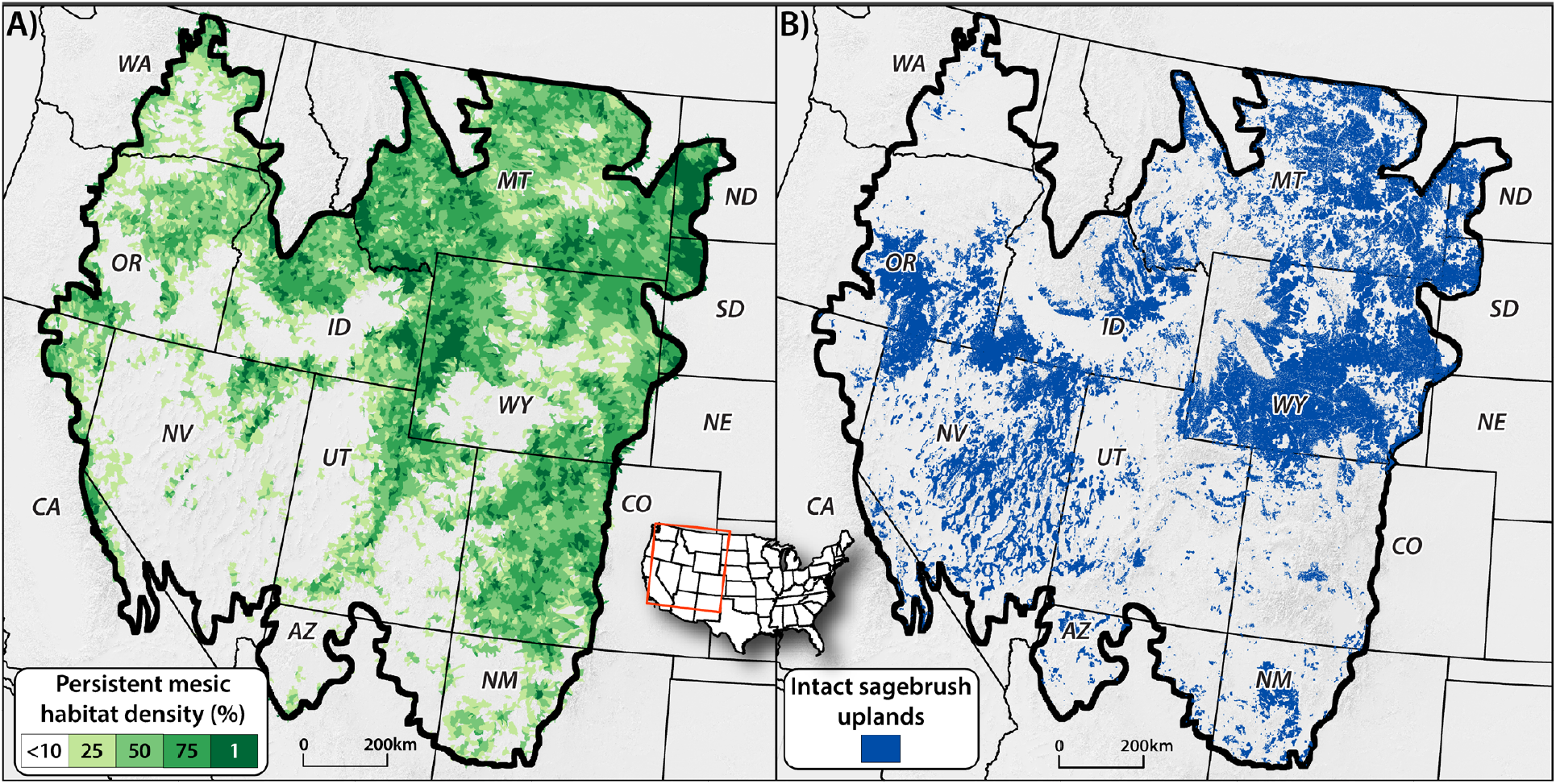
Maps showing A) the distribution of persistent mesic habitats, summarized within Hydrologic Unit Code 12 watersheds in the sagebrush biome, and B) Sage-brush Conservation Design, 2020, including intact uplands (Core Sagebrush Areas and Growth Opportunity Areas). Mesic habitat density indicates the proportion of the valley bottom area in a watershed with moderate or high vegetation persistence. Note the limited spatial overlap between areas of high mesic density (A) and intact sagebrush uplands (B), particularly evident in eastern Oregon and central Wyoming.

Regional patterns reveal distinct hydrologic and climatic drivers of mesic-upland separation (Fig. 3). We identify three distinct regions within the biome based on differences in climate and seasonal precipitation, using aggregated North American level III ecoregions (https://www.epa.gov/eco-research/ecoregions-north-america) (Smith et al., 1997). The Great Basin region was combined with the Columbia Plateau of eastern Washington, and the Rocky Mountains were combined with the Blue Mountains of northern Oregon and the Cascade/Sierra Mountains. These regional patterns dictate suitable restoration strategies, as floodplain functioning varies markedly across the sagebrush biome. The division between mesic habitats and sagebrush uplands manifests clearly at both watershed and valley bottom scales:

**Figure 3.**
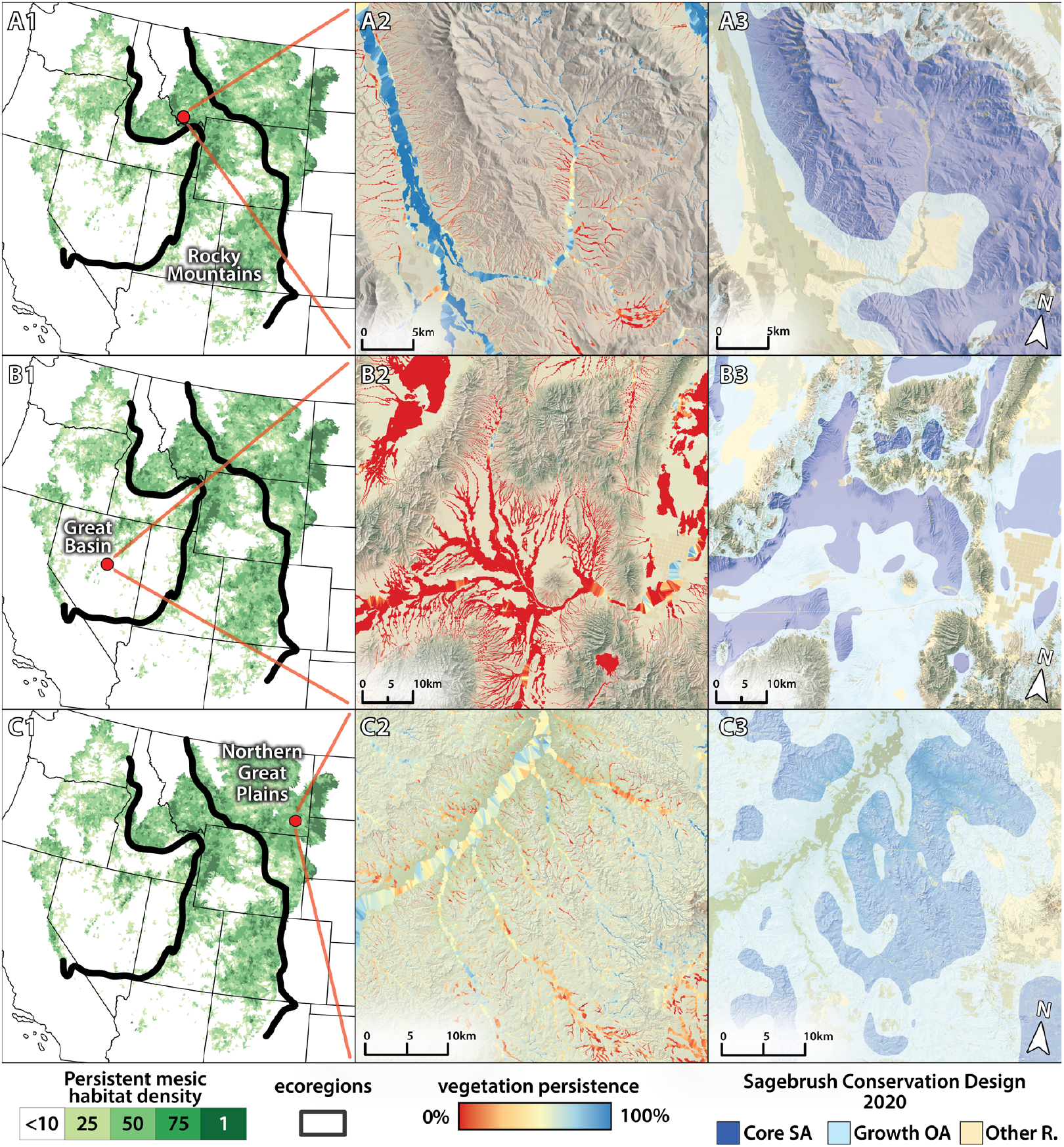
Regional patterns of vegetation persistence near intact sagebrush uplands. Rocky Mountains (A1-A3): highly persistent valley bottoms in mountain-foothill transition zones. Great Basin (B1-B3): limited persistence except near irrigation and mountain runoff. Northern Great Plains (C1-C3): mixed persistence patterns offering restoration opportunities.

#### Rocky Mountains

The highest overlap of persistent mesic vegetation and sagebrush uplands occurs where high-elevation forests transition into low-elevation rangelands (Fig. 3 A1-3). While this region contains the most persistent mesic vegetation, ecosystem overlap is limited because high-elevation forests dominate the landscape. Co-occurrence in the Rocky Mountains concentrates in foothill zones, where 13% of high-persistence mesic habitats lie within 5 kilometers of intact uplands, mostly in Northeast Idaho and Southwest Montana.

#### Great Basin

Here, water scarcity severely limits mesic vegetation persistence (Fig. 3 B1-3). The region’s cold winters and hot, arid summers prevent mesic vegetation from persisting without supplemental water (Naumburg et al., 2005). Persistent mesic areas are confined to irrigated lands or areas receiving sufficient runoff from sparse mountain ranges. Less than 6% of valley bottoms sustain high persistence, with only 3% of these limited mesic resources occurring near intact sagebrush uplands, underscoring severe water constraints on conservation potential.

#### Northern Great Plains

This region exhibits widespread co-occurrence of intact sagebrush uplands and moderate to high mesic persistence (Fig. 3 C1-3). Unlike the snowmelt-dependent ecosystems of the Rocky Mountains, the Northern Great Plains receive most of their precipitation during the summer months with highly variable rain events (Barker & Whitman, 1989). This summer moisture, though variable, sustains mesic vegetation more reliably than winter precipitation alone. Approximately 20% of high-persistence mesic habitats occur within 5 kilometers of intact sagebrush uplands in this region, with an additional 36% showing moderate persistence, revealing significant restoration opportunities where water availability is less limiting.

The limited spatial overlap between persistent mesic vegetation and intact sagebrush uplands reveals that conservation planning that only assesses upland vegetation condition is missing vital components of the sagebrush ecosystem. While this separation poses challenges for creating functional and connected landscapes, it also identifies strategic intervention opportunities. Water availability drives restoration potential: broadly, the Rocky Mountains region offers opportunities to conserve existing mesic habitats and reconnect water within floodplains, the Great Basin requires careful site selection where limited water and conservation goals align, and the Northern Great Plains provide intermediate conditions where natural variability could aid restoration efforts. These regional differences demonstrate that effective mesic conservation targeting requires regionally tailored restoration strategies based on local hydrologic conditions.

## Bridging the Gap: A Tiered Framework for Integrated Conservation

Addressing the spatial disconnect we identified requires a strategic approach to mesic conservation. To meet this need, we created a framework that combines two ecological factors: the persistence of mesic vegetation (as an indicator of water availability and conservation or restoration potential; described in methods) and proximity to intact sagebrush uplands (CSAs and GOAs). This framework further helps guide practitioners from broad watershed-scale prioritization to specific reach-scale implementation.

### Translating Ecological Patterns into Management Priorities

We developed a tiered management framework that translates ecological conditions into clear conservation actions at the subwatershed (HUC12) scale. Our classification system (Fig. 4) uses a hierarchical decision tree that first identifies subwatersheds, hereby referred to as “watersheds”, with sufficient nearby upland habitat (>15% of valley bottoms within 5 km of CSAs or GOAs) to support wildlife populations (Donnelly et al., 2016). For these watersheds, we prescribe management actions based on mesic vegetation patterns: protect and maintain watersheds with abundant high-persistence vegetation, restore and enhance watersheds with mixed persistence indicating hydrologic potential, or strategically manage watersheds with limited but present mesic resources. This approach produces three actionable management tiers (Figs 5&6):

**Figure 4.**
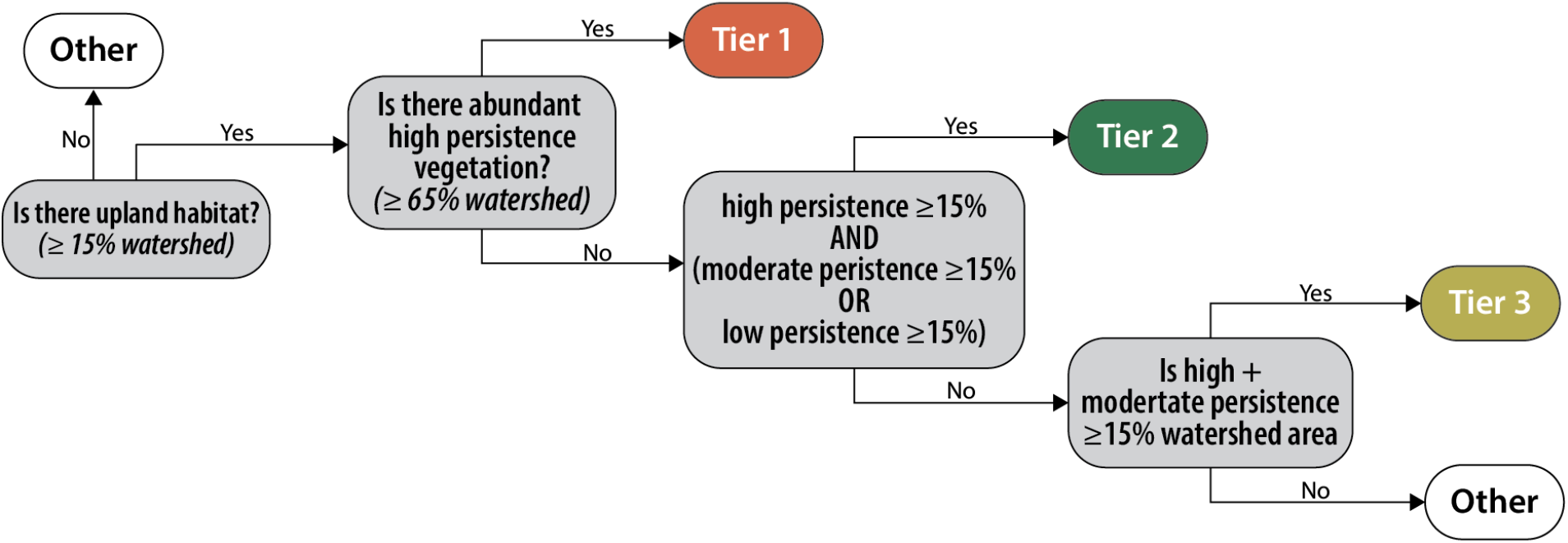
Decision tree for assigning management tiers to watersheds based on ecological conditions. Our framework’s decision-making process uses percentages that indicate the proportion of a watershed’s total valley bottom area. For example, ≥ 15% watershed means at least 15% of the watershed’s valley bottom area is near (within 5 km) Core Sagebrush Areas (CSAs) or Growth Opportunity Areas (GOAs).

**Figure 5.**
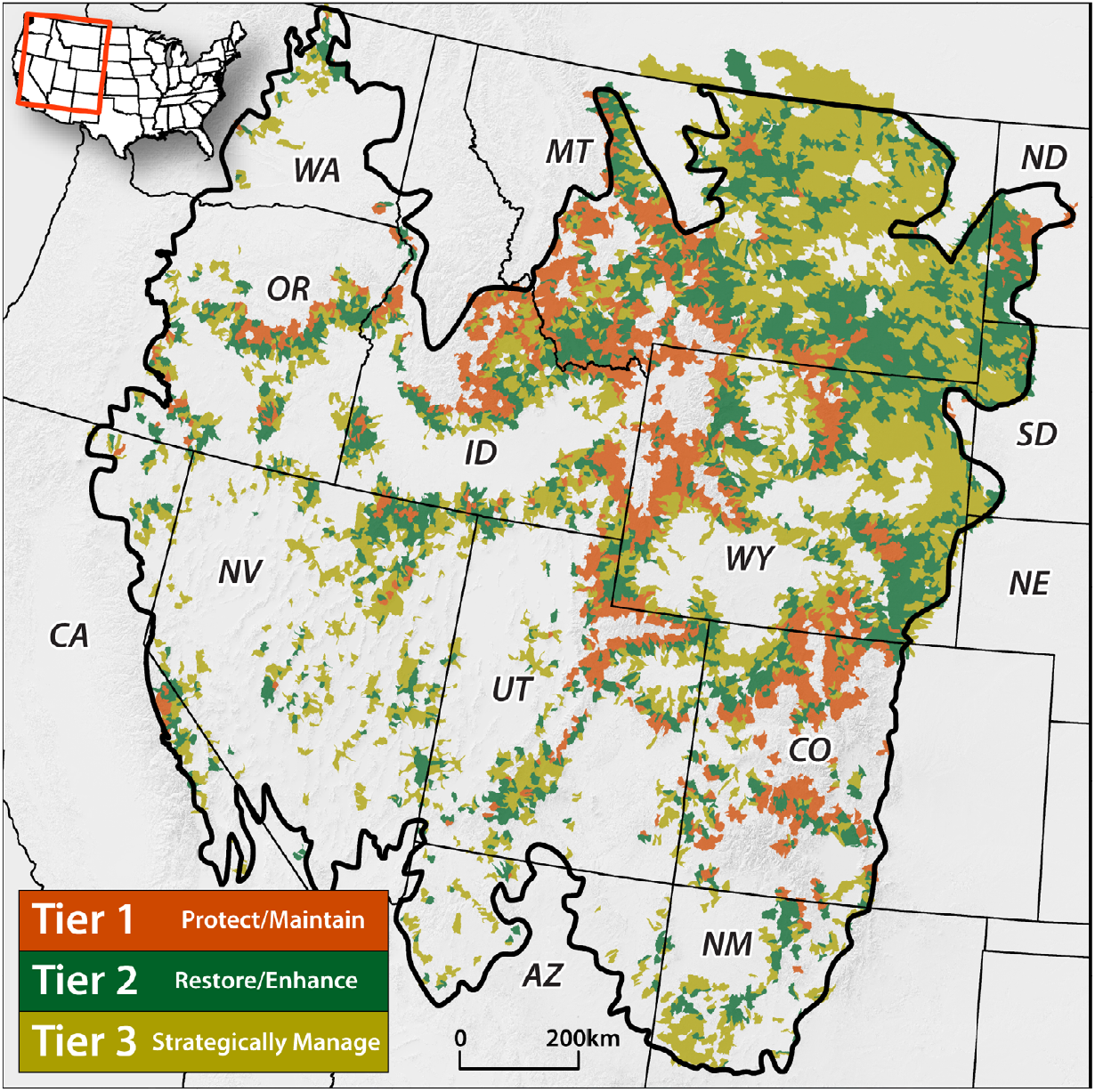
The framework classifies watersheds across the sagebrush biome based on management priorities for mesic valley bottoms near intact upland ecosystems. Tier 1 (red) watersheds: Protect and Maintain - high-functioning mesic valley bottoms requiring protection and maintenance. Tier 2 (orange) watersheds: Restore and Enhance - mixed persistence patterns indicating high restoration potential. Tier 3 (yellow) watersheds: Strategically Manage - limited mesic resources requiring targeted interventions in key areas.

#### Tier 1 - Protect and Maintain (9% of watersheds)

Focus protection and maintenance efforts in these high-value watersheds. These areas contain extensive high-persistence mesic habitats near intact sagebrush uplands, indicating hydrologically connected and stable mesic systems that require only targeted maintenance to sustain existing function.

#### Tier 2 - Restore and Enhance (14% of watersheds)

Prioritize restoration and enhancement in these watersheds offering the highest potential return on management investment. Valley bottoms here show mixed persistence patterns (high-persistence areas interspersed with moderate-persistence zones), suggesting water is present but not effectively distributed throughout all floodplains, making them ideal for intervention.

#### Tier 3 - Strategically Manage (17% of watersheds)

Apply strategic, targeted management in key areas to sustain or improve existing mesic habitats. These watersheds have limited but present mesic resources where selective interventions can maintain critical habitat features despite water limitations.

#### Other (60% of watersheds)

These watersheds represent alternative conservation strategies, as they either lack sufficient intact sagebrush upland habitat or have limited mesic resources that do not meet the tiered thresholds.

This tiered classification provides a practical framework for prioritizing mesic habitat conservation and restoration efforts across the sagebrush biome within interconnected upland sagebrush systems. However, effective implementation depends on social capacity, such as landowner willingness, collaborative networks, and supportive administrative conditions, as much as ecological factors (Wollstein et al., 2024). These social dimensions are particularly vital because, unlike many of the surrounding uplands, opportunities for mesic restoration concentrate on private lands. Specifically, Tier 1 watersheds are nearly evenly divided between public (54%) and private (46%) lands, while Tiers 2 and 3 contain predominantly privately owned valley bottoms (63% and 65%; Fig. 6B). While most valley bottoms in Tiers 2 and 3 are privately owned, a substantial portion of priority valley bottoms (in Tier 2-3)—ranging from 35% to 54%—still lie on public lands. This shared footprint underscores the need for cross-boundary, multi-jurisdictional partnerships. Because mesic systems and their associated hydrology operate independently of property lines, achieving ‘whole watershed’ restoration requires aligning public land management with voluntary private land conservation to ensure ecological benefits are realized across the entire landscape. This ownership pattern necessitates strong partnerships with agricultural producers and voluntary conservation programs. Within prioritized watersheds, practitioners can then target specific stream reaches where restoration is both ecologically sound and socially feasible.

**Figure 6.**
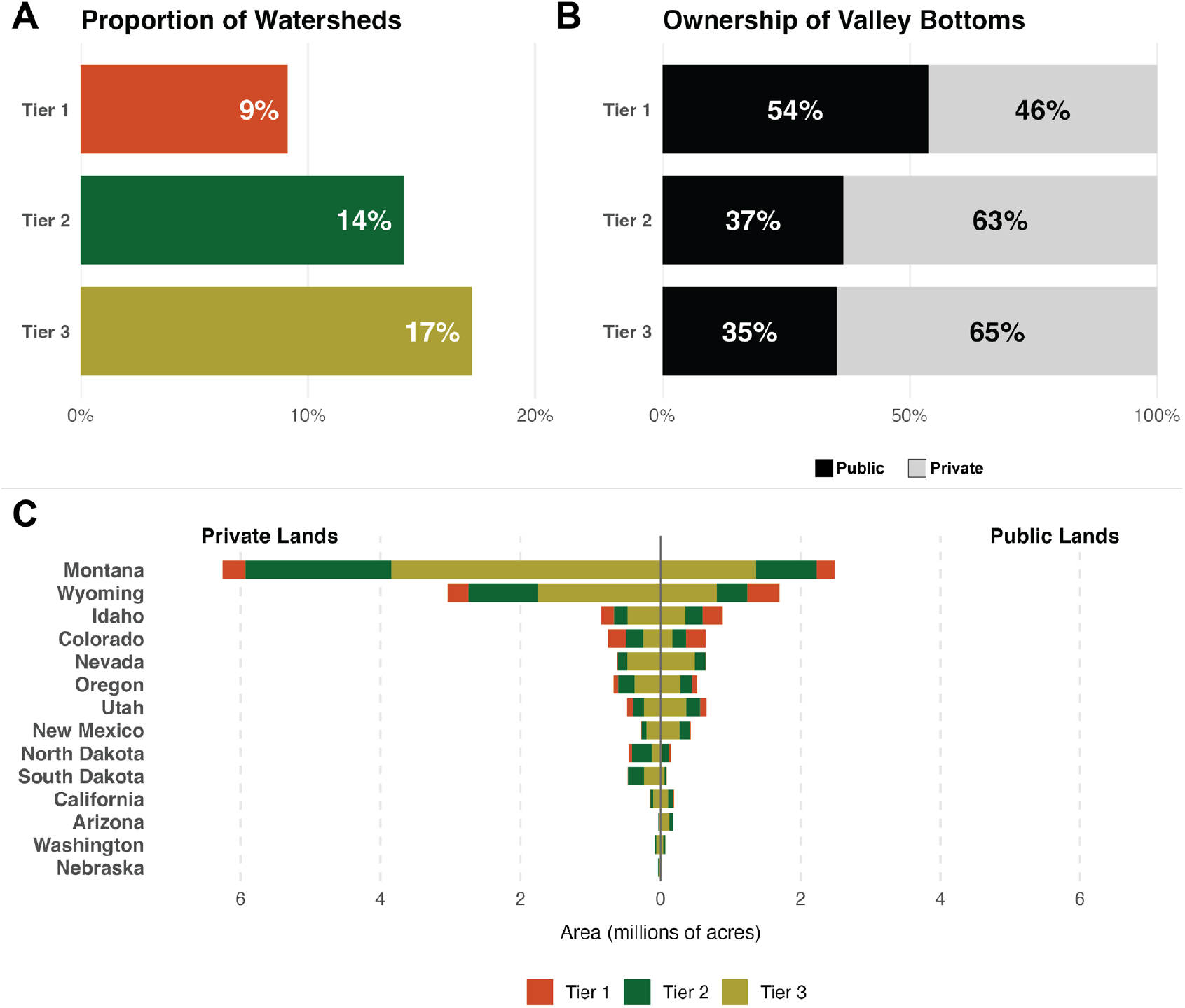
Characteristics of watershed tiers across the sagebrush biome, showing (A) the proportional distribution of all watersheds, (B) the composition of public versus private land ownership within valley bottoms, and (C) the total area of valley bottoms (in acres) within each state.

This restoration capacity also varies across the biome and by region and state (Fig. 6C). Consistent with the spatial distribution of mesic habitats (Fig. 3), watersheds in mountainous areas and those with more frequent precipitation events (Rocky Mountains and Northern Great Plains, respectively) yield varying densities of opportunity across the biome. For instance, the high density of Tier 2 and 3 watersheds in Montana provides a wealth of options but requires practitioners to apply additional social or logistical filters to decide where to work. Conversely, in the Great Basin, where priority zones are more distinct, the tiers serve as a surgical instrument for identifying high-impact areas, illustrating the trade-off between managing for landscape-scale abundance and targeting local resilience. Framework tier and ownership distribution figures are provided in the Supplemental Figures section.

### From Watersheds to Reaches: Implementation Guidance

Once practitioners identify priority watersheds, the next step is selecting specific valley bottom reaches for intervention. This shift to a finer spatial scale is crucial, as practical restoration techniques, like Low-Tech Process-Based Restoration (LTPBR), are implemented at the reach level, often in smaller stream floodplains where hydrologic processes are most readily modified (Wheaton et al., 2019). Our framework streamlines reach selection by leveraging persistence patterns along the stream network. We assess valley bottom reaches by comparing their vegetation persistence to conditions immediately upstream and downstream. Reaches with lower persistence bordered by areas of higher persistence may indicate localized hydrologic disconnection (Fig. 7A-B). These disconnected reaches indicate potential restoration sites for three reasons: 1) the presence of persistent vegetation above and below indicates water availability within the system; 2) the localized scale enables successful, targeted interventions; and 3) reconnection restores longitudinal connectivity throughout the stream network. For example, a reach with moderate persistence surrounded by high-persistence areas indicates that water exists in the system but is not effectively reaching the floodplain at that location. This pattern typically results from channel incision or other disturbances that separate the stream from its floodplain (Poff et al., 1997). LTPBR techniques can effectively treat these localized disconnections by dissipating stream energy, raising water tables, and reconnecting channels with their floodplains (Pollock et al., 2014; Zeedyk & Clothier, 2014).

**Figure 7.**
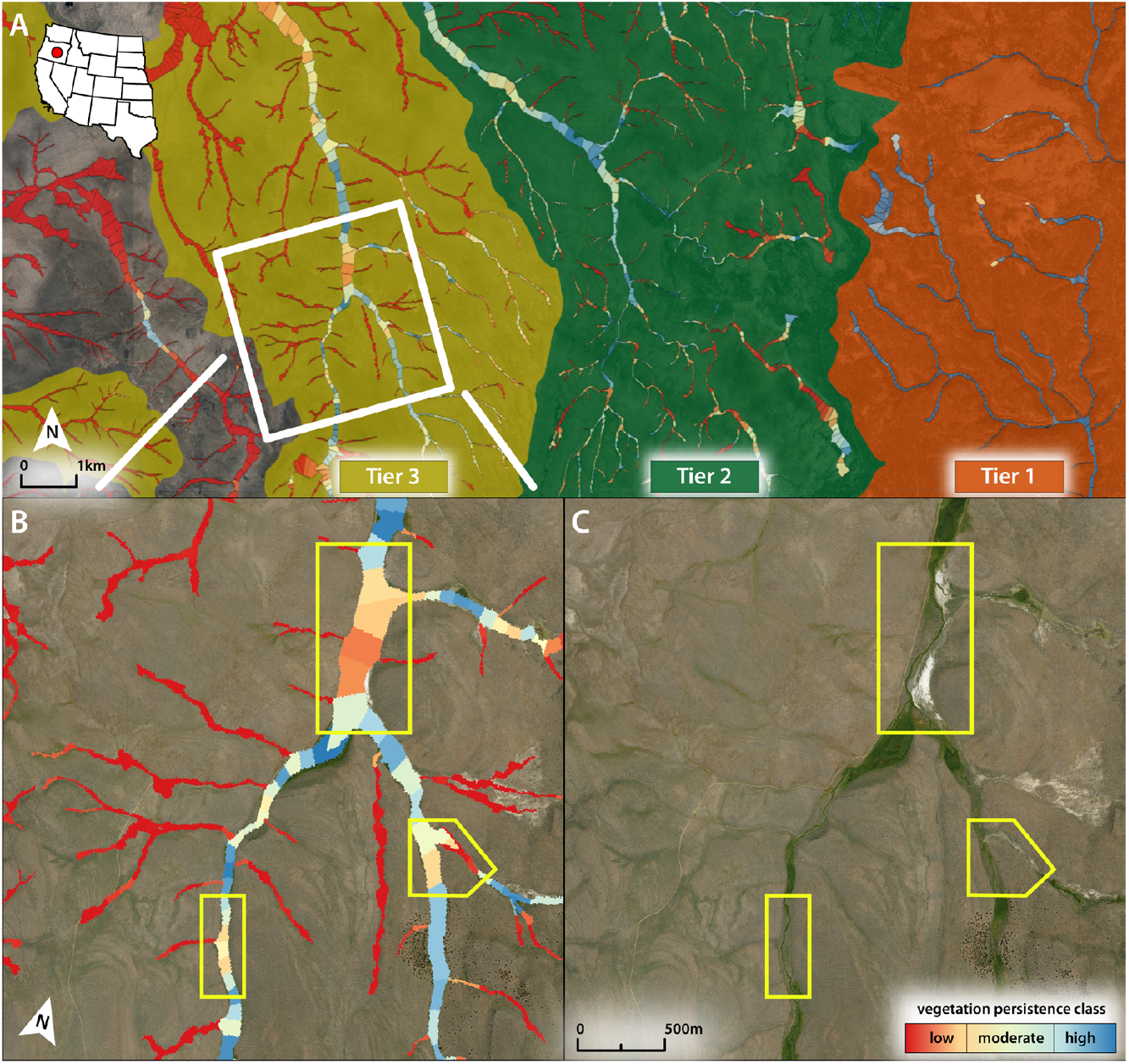
Potential restoration opportunities in a Tier 3 watershed in Central Oregon were identified using mesic vegetation persistence classes (panels A and B). Floodplain reaches with low to moderate persistence (outlined in yellow), bordered by high-persistence upstream and downstream reaches, indicate sustained water availability and represent priority restoration targets where localized interventions could reconnect the stream network. Persistence classes are derived from the mean 30m pixel values within reach segments. Panel C shows these reaches within a Tier 3 watershed, near Tier 1 and 2 watersheds, containing higher-persistence valley bottoms.

This multi-scale approach enables managers to efficiently filter thousands of watersheds in the sagebrush biome down to specific valley bottom reaches where restoration investment is most likely to yield maximum returns (Fig. 7C). By combining broad-scale remote sensing patterns with field verification of local conditions, practitioners can strategically allocate limited restoration resources to achieve the greatest ecological benefits.

## Putting Science into Practice: The Mesic Analysis Platform

To put our prioritization framework directly into the hands of practitioners, we developed the Mesic Analysis Platform. This interactive, web-based tool allows land managers and conservation practitioners to work in a dynamic environment where they can identify key landscapes for mesic restoration and monitor long-term vegetation responses. The platform integrates mesic vegetation persistence with data on intact sagebrush uplands, land ownership, vegetation composition, and other contextual landscape attributes, allowing users to apply our framework across multiple spatial scales. The platform supports three core functions for mesic conservation planning:

Watershed Prioritization (HUC 12): Users can apply our identified three-tier framework (Fig. 5) or develop custom strategies tailored to specific conservation objectives. The co-occurrence mapping feature dynamically identifies watersheds with the highest co-occurrence of user-defined variables (e.g., intact uplands, high-persistence mesic vegetation, or low-persistence vegetation), supporting customized, data-driven prioritization.

Stream Reach Targeting: The reach-summarized data layer enables users to identify specific valley bottom reaches suitable for restoration. Following our framework (Section 3), reaches with lower persistence flanked by higher persistence upstream and downstream indicate hydrologic disconnection and become priority targets for intervention (Fig. 7). Additional metrics, including tree cover, valley bottom characteristics, and flow type, further inform site selection.

Vegetation and Climate Trends: The platform provides a 41-year time series of key ecological data, including vegetation productivity (mean late-season NDVI), vegetation composition (Rangeland Analysis Platform), and climate data (GRIDMET precipitation and drought indices) for any selected reach, point, or user-defined polygon in the biome. This historical record enables users to monitor trends, evaluate previous restoration efforts, and assess mesic vegetation resilience to climate variability and management actions.

By integrating multi-scale data layers and analytical functions, the Mesic Analysis Platform bridges the gap between landscape-scale planning and on-the-ground implementation. Users can progress from identifying priority watersheds to selecting specific restoration sites while leveraging historical data for adaptive management and long-term project evaluation.

## A New Perspective for a Resilient Sagebrush Biome

Our analysis reveals a critical disconnect across western U.S. rangelands: highly persistent mesic vegetation rarely co-occurs with intact sagebrush ecosystems (Table 1). While Tier 1 watersheds containing both abundant mesic habitats and intact upland sagebrush systems comprise only 9% of the biome, significant restoration opportunities exist in Tier 2 (14%) and Tier 3 (17%) watersheds (Fig. 6A). Crucially, broad-scale mesic restoration depends on private working lands, which contain 63-65% of the valley bottoms in restoration-priority watersheds (Fig. 6B). This finding confirms that integrated sagebrush-mesic conservation requires addressing both the ecological disconnect and the social and administrative factors governing restoration on private lands. Success demands voluntary incentives, technical guidance, and strong partnerships with agricultural producers to restore degraded riparian areas and reconnect fragmented habitats (Sweeney & Blaine, 2016).

As with any framework based on remote sensing, successful implementation requires local expertise and validation. The 30-meter spatial resolution of the satellites cannot fully capture small stream dynamics or localized hydrological conditions that ultimately determine restoration feasibility. Furthermore, while we map watersheds where uplands and persistent mesic vegetation co-occur, mesic habitats outside these mapped areas may serve other localized conservation objectives. Ground validation and local knowledge of water rights, land use patterns, and site-specific conditions are essential to verify the restoration potential of candidate reaches identified through our data products.

The benefits of this integrated framework are bidirectional, informing both mesic restoration and upland conservation strategies. Just as we use SCD classifications to target mesic restoration that supports sagebrush conservation, our tier framework can reciprocally inform upland conservation priorities. Focusing sagebrush restoration efforts near Tier 1 watersheds ensures that investments in sagebrush habitat are supported by high-quality, mesic habitats. This reciprocal approach recognizes that wildlife require both habitat types in proximity, confirming that conservation succeeds when it addresses the complementary functions of the entire ecosystem (Donnelly et al., 2018; Thomas et al., 1979). To translate these findings into actions for a resilience sagebrush biome, we recommend that practitioners adopt the following approaches:

1. Restoration of mesic habitats should prioritize Tier 2 and 3 watersheds by focusing project funding and technical assistance where mesic restoration potential aligns with intact sagebrush ecosystems.
2. Engage private landowners where the broadest restoration opportunities occur across the biome to scale up voluntary, incentive-based conservation programs.
3. Adopt reciprocal conservation strategies by targeting degraded uplands near Tier 1 watersheds and intact mesic habitats, ensuring investments benefit the entire ecosystem.
4. Foster multi-jurisdictional partnerships to align public and private management, achieving “whole watershed” mesic conservation planning that transcends property lines.

## Data and Code Availability

The Mesic Analysis Platform (https://wlfw-um.projects.earthengine.app/view/mesic-analysis-platform) hosts the framework tiers and valley bottom reach extents for public interactive viewing and downloading. Data downloads are facilitated by either state or county outline within the sagebrush biome. Source code regarding data generation, tables, and figures has been permanently archived in a Zenodo repository (https://doi.org/10.5281/zenodo.17904662). Google Earth Engine and R scripts required for reproducibility are accessible via a GitHub repository (https://github.com/krisMueller/Beyond_Uplands_Mesic_Frameworks).

## Acknowledgements

This research used compute and storage resources provided by Google Earth Engine. Funding for this research was provided by the US Bureau of Land Management of Montana/Dakotas (grant code: F21AC00546) through the Intermountain West Joint Venture (IWJV) with help from the USDA-NRCS Working Lands for Wildlife.

## Author Contributions

K.R.M. led the analysis methodology, MAP application development, and wrote the initial draft. S.L.M. provided technical support for analysis methods. J.D.M. and D.E.N. assisted in analysis, summary decisions, and practical outcomes of the results and applications. All authors contributed feedback and comments during the writing process and have approved the final revision.

## References

Allred, B. W., Bestelmeyer, B. T., Boyd, C. S., Brown, C., Davies, K. W., Duniway, M. C., Ellsworth, L. M., Erickson, T. A., Fuhlendorf, S. D., Griffiths, T. V., Jansen, V., Jones, M. O., Karl, J., Knight, A., Maestas, J. D., Maynard, J. J., McCord, S. E., Naugle, D. E., Starns, H. D., … Uden, D. R. (2021). Improving Landsat predictions of rangeland fractional cover with multitask learning and uncertainty. Methods in Ecology and Evolution, 12(5), 841–849. 10.1111/2041-210X.13564

Barker, W. T., & Whitman, W. C. (1988). Vegetation of the Northern Great Plains. Rangelands, 10(6), 266–272.

Doherty, K., Theobald, D. M., Bradford, J. B., Wiechman, L. A., Bedrosian, G., Boyd, C. S., Cahill, M., Coates, P. S., Creutzburg, M. K., Crist, M. R., Finn, S. P., Kumar, A. V., Littlefield, C. E., Maestas, J. D., Prentice, K. L., Prochazka, B. G., Remington, T. E., Sparklin, W. D., Tull, J. C., … Zeller, K. A. (2022). A sagebrush conservation design to proactively restore America’s sagebrush biome. In Open-File Report (Nos. 2022–1081). U.S. Geological Survey. 10.3133/ofr20221081

Donnelly, J. P., Allred, B. W., Perret, D., Silverman, N. L., Tack, J. D., Dreitz, V. J., Maestas, J. D., & Naugle, D. E. (2018). Seasonal drought in North America’s sagebrush biome structures dynamic mesic resources for sage-grouse. Ecology and Evolution, 8(24), 12492–12505. 10.1002/ece3.4614

Donnelly, J. P., Collins, D. P., Knetter, J. M., Gammonley, J. H., Boggie, M. A., Grisham, B. A., Nowak, M. C., & Naugle, D. E. (2024). Flood-irrigated agriculture mediates climate-induced wetland scarcity for summering sandhill cranes in western North America. Ecology and Evolution, 14(3), e10998. 10.1002/ece3.10998

Donnelly, J. P., Naugle, D. E., Hagen, C. A., & Maestas, J. D. (2016). Public lands and private waters: Scarce mesic resources structure land tenure and sage-grouse distributions. Ecosphere, 7(1), e01208. 10.1002/ecs2.1208

Dudgeon, D., Arthington, A. H., Gessner, M. O., Kawabata, Z.-I., Knowler, D. J., Lévêque, C., Naiman, R. J., Prieur-Richard, A.-H., Soto, D., Stiassny, M. L. J., & Sullivan, C. A. (2006). Freshwater biodiversity: Importance, threats, status and conservation challenges. Biological Reviews of the Cambridge Philosophical Society, 81(2), 163–182. 10.1017/S1464793105006950

Gilbert, J. T., Macfarlane, W. W., & Wheaton, J. M. (2016). The Valley Bottom Extraction Tool (V-BET): A GIS tool for delineating valley bottoms across entire drainage networks. Computers & Geosciences, 97, 1–14. 10.1016/j.cageo.2016.07.014

Gorelick, N., Hancher, M., Dixon, M., Ilyushchenko, S., Thau, D., & Moore, R. (2017). Google Earth Engine: Planetary-scale geospatial analysis for everyone. Remote Sensing of Environment. https://earthengine.google.com/faq/

Kolarik, N. E., Roopsind, A., Pickens, A., & Brandt, J. S. (2023). A satellite-based monitoring system for quantifying surface water and mesic vegetation dynamics in a semi-arid region. Ecological Indicators, 147, 109965. 10.1016/j.ecolind.2023.109965

Lundblad, C. G., Hagen, C. A., Donnelly, J. P., Vold, S. T., Moser, A. M., & Espinosa, S. P. (2022). Sensitivity to weather drives Great Basin mesic resources and Greater Sage-Grouse productivity. Ecological Indicators, 142, 109231. 10.1016/j.ecolind.2022.109231

Maestas, J. D., Wheaton, J. M., Bouwes, N., Swanson, S. R., & Dickard, M. (2023). Water Is Life: Importance and Management of Riparian Areas for Rangeland Wildlife. In L. B. McNew, D. K. Dahlgren, & J. L. Beck (Eds.), Rangeland Wildlife Ecology and Conservation (pp. 177–208). Springer International Publishing. 10.1007/978-3-031-34037-6_7

Manning, A., Julian, J. P., & Doyle, M. W. (2020). Riparian vegetation as an indicator of stream channel presence and connectivity in arid environments. Journal of Arid Environments, 178, 104167. 10.1016/j.jaridenv.2020.104167

Mozelewski, T. G., Freeman, P. T., Kumar, A. V., Naugle, D. E., Olimpi, E. M., Morford, S. L., Jeffries, M. I., Pilliod, D. S., Littlefield, C. E., McCord, S. E., Wiechman, L. A., Kachergis, E. J., & Doherty, K. E. (2024). Closing the Conservation Gap: Spatial Targeting and Coordination are Needed for Conservation to Keep Pace with Sagebrush Losses. Rangeland Ecology & Management, 97, 12–24. 10.1016/j.rama.2024.08.016

Naiman, R. J., Decamps, H., & Pollock, M. (1993). The Role of Riparian Corridors in Maintaining Regional Biodiversity. Ecological Applications, 3(2), 209–212. 10.2307/1941822

Naumburg, E., Mata-gonzalez, R., Hunter, R. G., Mclendon, T., & Martin, D. W. (2005). Phreatophytic Vegetation and Groundwater Fluctuations: A Review of Current Research and Application of Ecosystem Response Modeling with an Emphasis on Great Basin Vegetation. Environmental Management, 35(6), 726–740. 10.1007/s00267-004-0194-7

Nilsson, C., & Berggren, K. (2000). Alterations of Riparian Ecosystems Caused by River Regulation: Dam operations have caused global-scale ecological changes in riparian ecosystems. How to protect river environments and human needs of rivers remains one of the most important questions of our time. BioScience, 50(9), 783–792. 10.1641/0006-3568(2000)050[0783:AORECB]2.0.CO;2

Noy-Meir, I. (1973). Desert Ecosystems: Environment and Producers. Annual Review of Ecology, Evolution and Systematics, 4(Volume 4, 1973), 25–51. 10.1146/annurev.es.04.110173.000325

Patten, D. T. (1998). Riparian ecosytems of semi-arid North America: Diversity and human impacts. Wetlands, 18(4), 498–512. 10.1007/BF03161668

Poff, N. L., Allan, J. D., Bain, M. B., Karr, J. R., Prestegaard, K. L., Richter, B. D., Sparks, R. E., & Stromberg, J. C. (1997). The Natural Flow Regime. BioScience, 47(11), 769–784. 10.2307/1313099

Pollock, M. M., Beechie, T. J., Wheaton, J. M., Jordan, C. E., Bouwes, N., Weber, N., & Volk, C. (2014). Using Beaver Dams to Restore Incised Stream Ecosystems. BioScience, 64(4), 279–290. 10.1093/biosci/biu036

Riis, T., Kelly-Quinn, M., Aguiar, F. C., Manolaki, P., Bruno, D., Bejarano, M. D., Clerici, N., Fernandes, M. R., Franco, J. C., Pettit, N., Portela, A. P., Tammeorg, O., Tammeorg, P., Rodríguez-González, P. M., & Dufour, S. (2020). Global Overview of Ecosystem Services Provided by Riparian Vegetation. BioScience, 70(6), 501–514. 10.1093/biosci/biaa041

Rodríguez-González, P. M., Abraham, E., Aguiar, F., Andreoli, A., Baležentienė, L., Berisha, N., Bernez, I., Bruen, M., Bruno, D., Camporeale, C., Čarni, A., Chilikova-Lubomirova, M., Corenblit, D., Ćušterevska, R., Doody, T., England, J., Evette, A., Francis, R., Garófano-Gómez, V., … Dufour, S. (2022). Bringing the margin to the focus: 10 challenges for riparian vegetation science and management. WIREs Water, 9(5), e1604. 10.1002/wat2.1604

Schneider, C., Flörke, M., De Stefano, L., & Petersen-Perlman, J. D. (2017). Hydrological threats to riparian wetlands of international importance – a global quantitative and qualitative analysis. Hydrology and Earth System Sciences, 21(6), 2799–2815. 10.5194/hess-21-2799-2017

Shrestha, N., Kolarik, N. E., & Brandt, J. S. (2024). Mesic vegetation persistence: A new approach for monitoring spatial and temporal changes in water availability in dryland regions using cloud computing and the sentinel and Landsat constellations. Science of The Total Environment, 917, 170491. 10.1016/j.scitotenv.2024.170491

Silverman, N. L., Allred, B. W., Donnelly, J. P., Chapman, T. B., Maestas, J. D., Wheaton, J. M., White, J., & Naugle, D. E. (2019). Low-tech riparian and wet meadow restoration increases vegetation productivity and resilience across semiarid rangelands. Restoration Ecology, 27(2), 269–278. 10.1111/rec.12869

Smith, T. M., Shugart, H. H., & Woodward, F. I. (1997). Plant Functional Types: Their Relevance to Ecosystem Properties and Global Change. Cambridge University Press.

Sweeney, B. W., & Blaine, J. G. (2016). River conservation, restoration, and preservation: Rewarding private behavior to enhance the commons. Freshwater Science, 35(3), 755–763. 10.1086/687364

Theobald, D. M., Merritt, D. M., & Norman, J. B. (2010). Assessment of threats to riparian ecosystems in the western US. Fort Collins, CO, USA: Colorado State University.

Thomas, J. W., Maser, C., & Rodiek, J. E. (1979). Wildlife Habitats in Managed Rangelands: The Great Basin of Southeastern Oregon : Riparian Zones. Pacific Northwest Forest and Range Experiment Station, U.S. Department of Agriculture, Forest Service.

U.S. Geological Survey (USGS) Gap Analysis Project (GAP), 2024, Protected Areas Database of the United States (PAD-US) 4.1: U.S. Geological Survey data release, 10.5066/P96WBCHS.

Weier, J., & Herring, D. (2000, August 30). Measuring Vegetation (NDVI & EVI) [Text.Article]. NASA Earth Observatory. https://earthobservatory.nasa.gov/features/MeasuringVegetation

Wheaton, J. M., Bennett, S. N., Bouwes, N., Camp, R., Maestas, J. D., & Shahverdian, S. (2019). Chapter 2 – Principles of Low-Tech Process-Based Restoration. 10.13140/RG.2.2.34270.69447

Wollstein, K., Johnson, D., & Boyd, C. (2024). Assessing Conservation Readiness: The Where, Who, and How of Strategic Conservation in the Sagebrush Biome. Rangeland Ecology & Management, 97, 187–199. 10.1016/j.rama.2024.08.013

Zeedyk, B., & Clothier, V. (2014). Let the Water Do the Work: Induced Meandering, an Evolving Method for Restoring. Chelsea Green Publishing.

